# TAK-071, a muscarinic M_1_ receptor positive allosteric modulator, attenuates scopolamine-induced quantitative electroencephalogram power spectral changes in cynomolgus monkeys

**DOI:** 10.1101/468686

**Authors:** Emi Kurimoto, Masato Nakashima, Haruhide Kimura, Motohisa Suzuki

**Affiliations:** Neuroscience Drug Discovery Unit, Research, Takeda Pharmaceutical Company Limited, Fujisawa, Kanagawa, Japan

## Abstract

Activation of the muscarinic M_1_ receptor is a promising approach to improve cognitive deficits associated with cholinergic dysfunction in Alzheimer’s disease, dementia with Lewy bodies, and schizophrenia. TAK-071 is an M_1_-selective positive allosteric modulator that improves cognitive deficits induced by scopolamine, a non-selective muscarinic receptor antagonist, with reduced side effects on gastrointestinal function in rats. In this study, we explored changes in quantitative electroencephalography (qEEG) power bands, with or without scopolamine challenge, as a non-invasive translational biomarker for the effect of TAK-071 in cynomolgus monkeys. Scopolamine has been reported to increase theta and delta power bands and decrease alpha power band in healthy volunteers. In line with the clinical observations, scopolamine (25–100 µg/kg, subcutaneous administration [s.c.]) increased theta and delta power bands in cynomolgus monkeys in a dose-dependent manner, whereas it had the opposite effect on alpha power band. The effects of TAK-071 on scopolamine (25 µg/kg, s.c.)-induced qEEG spectral changes were examined using an acetylcholinesterase inhibitor donepezil and a muscarinic M_1_/M_4_ receptor agonist xanomeline as comparative cholinomimetics. TAK-071 (0.3–3 mg/kg, oral administration [p.o.]), donepezil (3 mg/kg, p.o.), and xanomeline (1 mg/kg, s.c.) suppressed the scopolamine-induced increases in alpha, theta, and delta power bands. These results suggest that changes in specific qEEG power bands, in particular theta and delta power bands in the context of scopolamine challenge, could be used as translational biomarkers for the evaluation of TAK-071 in clinical studies.

## Introduction

Alzheimer’s disease (AD) and dementia with Lewy bodies (DLB) are progressive neurodegenerative disorders that profoundly impact cognitive function. A substantial amount of evidence has established that degeneration of cholinergic neurons contributes to cognitive deterioration in patients with AD and DLB [1-3]. The effects of currently available cholinergic treatments are mediated by increasing the synaptic acetylcholine (ACh) concentration through inhibition of acetylcholinesterase (AChE), leading to symptomatic cognitive improvement. However, these treatments have limited efficacy in part because of severe gastrointestinal (GI) side effects. One approach to selectively restore cholinergic function in the brain and enhance cognitive function is direct activation of the neuronal muscarinic M_1_ receptor (M_1_R), as several lines of evidence have demonstrated that M_1_R is highly expressed in the brain regions associated with cognition, and its activators are expected to improve cognitive deficits in AD [4-9]. However, the development of subtype-selective drugs that bind to the orthosteric site of M_1_R remains challenging because of high conservation across the muscarinic receptor subfamily. To achieve high selectivity against M_1_R, we focused on the development of an M_1_ positive allosteric modulator (PAM). In our previous study, we reported that even highly selective M_1_ PAMs caused diarrhea in mice through M_1_R activation and that an ileum contractile response upon treatment with M_1_ PAMs was positively correlated with their α-values, indices of binding cooperativity between ACh and M_1_R [10]. Furthermore, M_1_ PAMs with low α-values caused less diarrhea side effects [10]. Based on this report, we sought M_1_ PAMs with lower α-values, and identified TAK-071 (4-fluoro-2-[(3*S*,4*S*)-4-hydroxytetrahydro-2*H*-pyran-3-yl]-5-methyl-6-[4-(1*H*-pyrazol-1-yl)benzyl]-2,3-dihydro-1*H*-isoindol-1-one) with sufficient cognitive efficacy and milder GI side effects in both monotherapy and combination therapy with AChE inhibitors such as donepezil and rivastigmine [11].

The main goals of early-phase drug development are demonstration of substantial target engagement at doses that are safe and tolerable and determination of a therapeutic dose range. To facilitate drug development, translational research requires minimally invasive and quantifiable biomarkers that can be obtained in both preclinical and clinical studies. Positron emission tomography (PET) is commonly used for drug development of the central nervous system (CNS) as it allows assessment of ligand-target binding and its distribution in the brain. The use of PET imaging is limited by the requirement for a well-characterized PET radioligand suitable for demonstration of target engagement. Recently, direct radiolabeling of a selective M_1_R allosteric agonist, GSK1034702, was performed and PET measurements revealed good brain uptake of this agonist [12]. However, regardless of good brain penetration, it has not been clarified that the radioligand specifically binds to an allosteric site of M_1_R. Furthermore, although PET studies can evaluate drug-receptor interactions in terms of target occupancy, the pharmacodynamic (PD) effects are not measured [13]. Therefore, alternative approaches are needed to demonstrate pharmacokinetic-pharmacodynamic (PK/PD) relationships and thereby provide evidence for functional modulation of the pharmacological target of TAK-071.

Following the first observations by Berger of abnormal electroencephalogram (EEG) sequences in patients with AD [14, 15], there have been many studies on quantitative EEG (qEEG) in patients with AD. The hallmarks of qEEG abnormalities in patients with AD and DLB is slowing of rhythms in specific brain regions, characterized by increases in theta and delta activities and decreases in alpha and beta activities under relaxed sitting conditions [16-19]. Correlation between the degree of the qEEG abnormality and cognitive impairment have been noted [20-22]. Moreover, it has been reported that donepezil significantly decreased delta activity and increased alpha and beta activities in patients with AD, suggesting normalization of the qEEG abnormalities in AD [23].

In healthy subjects, scopolamine, a non-selective muscarinic receptor antagonist, commonly used for pharmacological modeling of the cholinergic deficit in AD [24], caused concomitant appearance of transient cognitive impairments and qEEG spectral changes that resemble those observed in patients with AD and DLB [25-27]. After scopolamine treatment, qEEG resting state studies revealed an increase in delta power band and decrease in alpha power band. Moreover, AChE inhibitors such as donepezil, rivastigmine, and tacrine decreased delta power band after treatment with scopolamine in freely moving rats [28, 29].

In this study, we investigated whether qEEG analysis could function as a PD translational biomarker for TAK-071 efficacy assessment in non-human primates. To determine the utility of qEEG analysis, we examined whether the effects of scopolamine on changes in qEEG spectra in monkeys were similar to those in healthy subjects or in patients with AD and DLB. In addition, we evaluated the effects of cholinergic drugs such as donepezil, xanomeline, and TAK-071 on scopolamine-induced qEEG spectral changes. Our results provide evidence that qEEG analysis under the scopolamine challenge paradigm might be a valuable translational biomarker for the evaluation of TAK-071 in clinical trials.

## Materials and methods

### Ethics Statement

Animal handling and the experimental protocols used in this research were in accordance with the guidelines of the Institutional Animal Care and Use Committee of Takeda Pharmaceutical Company Limited (Protocol number: 00020938). Animal research facilities in Takeda Pharmaceutical Company Limited (Kanagawa, Japan) are accredited by the Association for Assessment and Accreditation of Laboratory Animal Care (AAALAC).

### Animals

Animal welfare protocols and the steps taken to ameliorate suffering were in accordance with the recommendations of the Weatherall report on the use of non-human primates in research. Male cynomolgus monkeys (*Macaca fascicularis*) were purchased from Hamri Company Limited (Ibaraki, Japan). The monkeys were individually housed and maintained in accordance with the guidelines approved by AAALAC. They were given water *ad libitum* and fed once daily with a complete, nutritionally balanced diet enriched with fruits and gummy candies. All monkeys, housed and handled in strict accordance with the guidelines for good animal practice under the supervision of veterinarians, received environmental enrichment and were monitored for evidence of disease and changes in attitude, appetite, or behavior suggestive of illness. In addition, every effort was made to alleviate animal discomfort and pain by routine use of appropriate anesthetic and analgesic agents.

### Surgical Procedures

Implant surgery was carried out at Hamri Company Limited (Ibaraki, Japan). Briefly, under isoflurane anesthesia (1–5%, Pfizer Japan Inc., Tokyo, Japan), male cynomolgus monkeys (3–5 years old, Hamri Co., Ltd., Ibaraki, Japan) were surgically implanted with radio-telemetry transmitters (TL10M3-D70-EEE, Data Sciences International Inc., St. Paul, MN). EEG leads were stereotactically positioned in the parietal area and secured to the cranium with stainless-steel screws in contact with the dura. Unilateral electrooculography (EOG) leads were positioned at the superior orbital margin of one eye and secured with stainless-steel screws. Bilateral electromyography (EMG) leads were implanted in the back cervical muscles. After surgery, each monkey was administered penicillin (100,000 units/animal, intramuscularly [i.m.], Meiji Seika Pharma Co., Ltd., Tokyo, Japan), buprenorphine (0.02 mg/kg, i.m., Otsuka Pharmaceutical Co., Ltd., Tokyo, Japan), and prednisolone (1 mg/kg, subcutaneously [s.c.], Kyoritsu Seiyaku Co., Ltd., Tokyo, Japan) daily for one week.

### Chemical treatment

TAK-071, 4-fluoro-2-[(3*S*,4*S*)-4-hydroxytetrahydro-2*H*-pyran-3-yl]-5-methyl-6-[4-(1*H*-pyrazol-1-yl)benzyl]-2,3-dihydro-1*H*-isoindol-1-one, was synthesized by Takeda Pharmaceutical Company Limited (Kanagawa, Japan). Donepezil hydrochloride and xanomeline oxalate were purchased from Mega Fine Pharma (P) Limited (Mumbai, India) and Metina AB (Lund, Sweden), respectively. Scopolamine hydrobromide was purchased from Tocris Bioscience (Ellisville, MO). TAK-071 and donepezil hydrochloride were suspended in 0.5% (w/v) methylcellulose in distilled water and the volume of administration was 2 mL/kg orally (p.o.). Xanomeline oxalate and scopolamine hydrobromide were dissolved in saline, and the volume of administration for xanomeline and scopolamine were 0.5 and 0.1 mL/kg (s.c.), respectively. The doses of compounds correspond to their salt forms. The administration was scheduled according to a 3×3 cross-over design.

### Measurements

After at least a 1-month recovery period in their home cages, the monkeys were habituated to the recording chamber (acrylic cage, 60 × 55 × 75 [cm^3^ width × depth × height]) located in a soundproof, electrically shielded room. Cortical EEG, EMG, and EOG signals were recorded at a sampling rate of 500 Hz using Dataquest ART software (Data Sciences International, New Brighton, MN). The behavioral and postural changes of the animals were observed using a video camera continuously throughout the experiment. The recordings were performed between 9 a.m. and 4 p.m..

EEG data were analyzed off-line using SleepSign (KISSEI COMTEC Co., Ltd., Nagano, Japan). Digitized EEG signals were high-pass filtered with a 0.75-Hz filter. Spectral power densities for each frequency band were calculated by fast Fourier transform in consecutive 10-s epochs. Epochs containing abnormal EEG, defined by excessive amplitude (> 500 µV), were omitted from data analysis. For analysis of temporal changes in a specific spectral band (1–4 Hz for delta, 4–8 Hz for theta, 8–12 Hz for alpha, 12–30 Hz for beta, 30–80 Hz for gamma), the EEG power of each spectral band was calculated every 15 min. For each animal, the qEEG spectral analyses were performed from 30 to 240 min after treatment because oral and subcutaneous administration of drugs to monkeys can affect the qEEG power spectra. Data obtained in the post-drug period were normalized with those obtained in the baseline period and were calculated as the area under the curve (AUC) between 30 and 210 min after drug administration for quantitative analyses.

### Statistical analysis

Data were expressed as mean ± standard error of the mean (SEM) and were analyzed using paired *t*-test, in which *P* ≤ 0.05 was considered significant. All analyses were performed using the SAS system 8 (SAS Institute, Cary, NC).

## Results

### Scopolamine increased alpha, theta, and delta power bands, in a dose-dependent manner

In clinical trials, multiple lines of evidence have suggested that scopolamine treatment induces qEEG power spectral changes, particularly increases in theta and delta power bands and decrease in alpha power band [27, 30, 31]. Initially, we examined the effects of scopolamine in qEEG power spectra in cynomolgus monkeys. To evaluate dose-dependency, scopolamine was subcutaneously administered at 25, 50, or 100 µg/kg to monkeys. Indeed, scopolamine increased alpha, theta, and delta power bands in a dose-dependent manner (Fig 1A–C). These effects in all power bands persisted for 240 min after treatment, and peak increases were observed between 60 and 90 min after treatment. The time-dependent changes of each qEEG spectrum during scopolamine treatment relatively corresponded to its PK profile (S1 Fig and S1 Table). Alpha, theta, and delta power bands were significantly increased at 50 and 100 µg/kg doses (*P* ≤ 0.05, Fig 1A–C). In contrast, beta and gamma power bands did not change remarkably in the dose range of 25–100 µg/kg of scopolamine (S2 Fig). It is reported that scopolamine at 20–50 µg/kg moderately caused cognitive impairment in monkeys [32, 33]. Therefore, the dose of 25 = µg/kg scopolamine was selected for further examinations.

**Fig 1.**
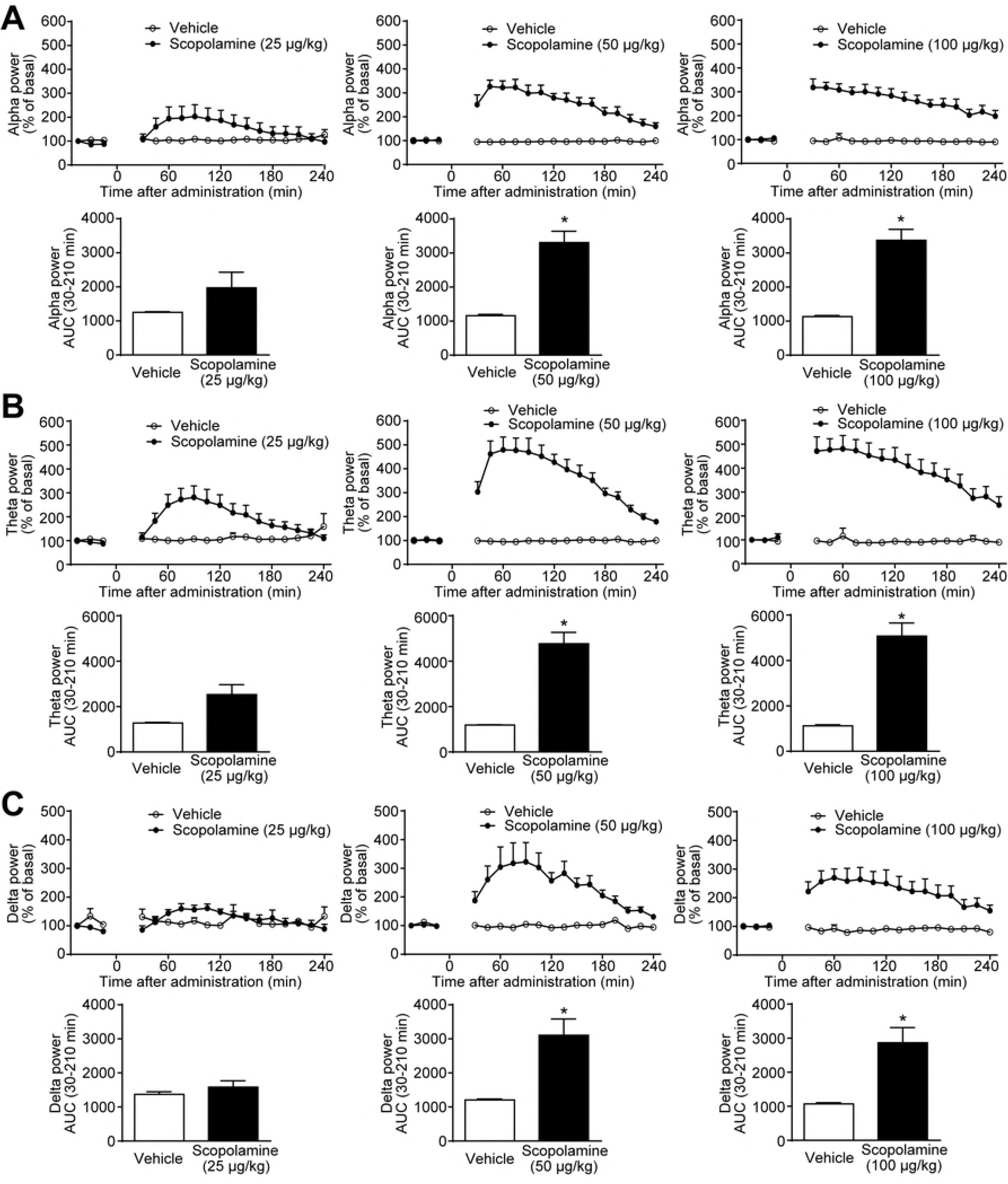
Scopolamine caused increases in alpha, theta, and delta power bands of the qEEG spectra, in a dose-dependent manner. Scopolamine (25–100 µg/kg) was subcutaneously administered to cynomolgus monkeys. After treatment with scopolamine or vehicle, (A) alpha, (B) theta, and (C) delta power bands of qEEG spectra were measured from 30 to 240 min. Results represent mean ± SEM of AUC between 30 and 210 min after treatment for 4 monkeys in each group. ^*^*P* ≤ 0.05 versus vehicle-treated group by paired *t*-test. AUC, area under the curve; qEEG, quantitative electroencephalogram; SEM, standard error of the mean.

### Donepezil, an AChE inhibitor, suppressed the scopolamine-induced qEEG spectral changes

We hypothesized that the scopolamine-induced qEEG spectral changes in monkeys would be counteracted by an AChE inhibitor donepezil through enhancement of ACh concentration in the synaptic cleft. Donepezil (3 mg/kg, p.o.) and scopolamine (25 µg/kg, s.c.) were simultaneously administered to monkeys. The dose of donepezil for this study was determined based on the previous report, which showed that oral treatment of donepezil at 1–3 mg/kg significantly attenuated the scopolamine-induced cognitive deficits in monkeys [33]. The increases in alpha and theta power bands induced by scopolamine treatment tended to be attenuated by donepezil (Fig 2A and B). Furthermore, in accordance with the previous report in rats [28], donepezil significantly suppressed the scopolamine-induced increase in the delta power band (Fig 2C). These effects were observed between 60 and 240 min after drug treatment (Fig 2A–C).

**Fig 2.**
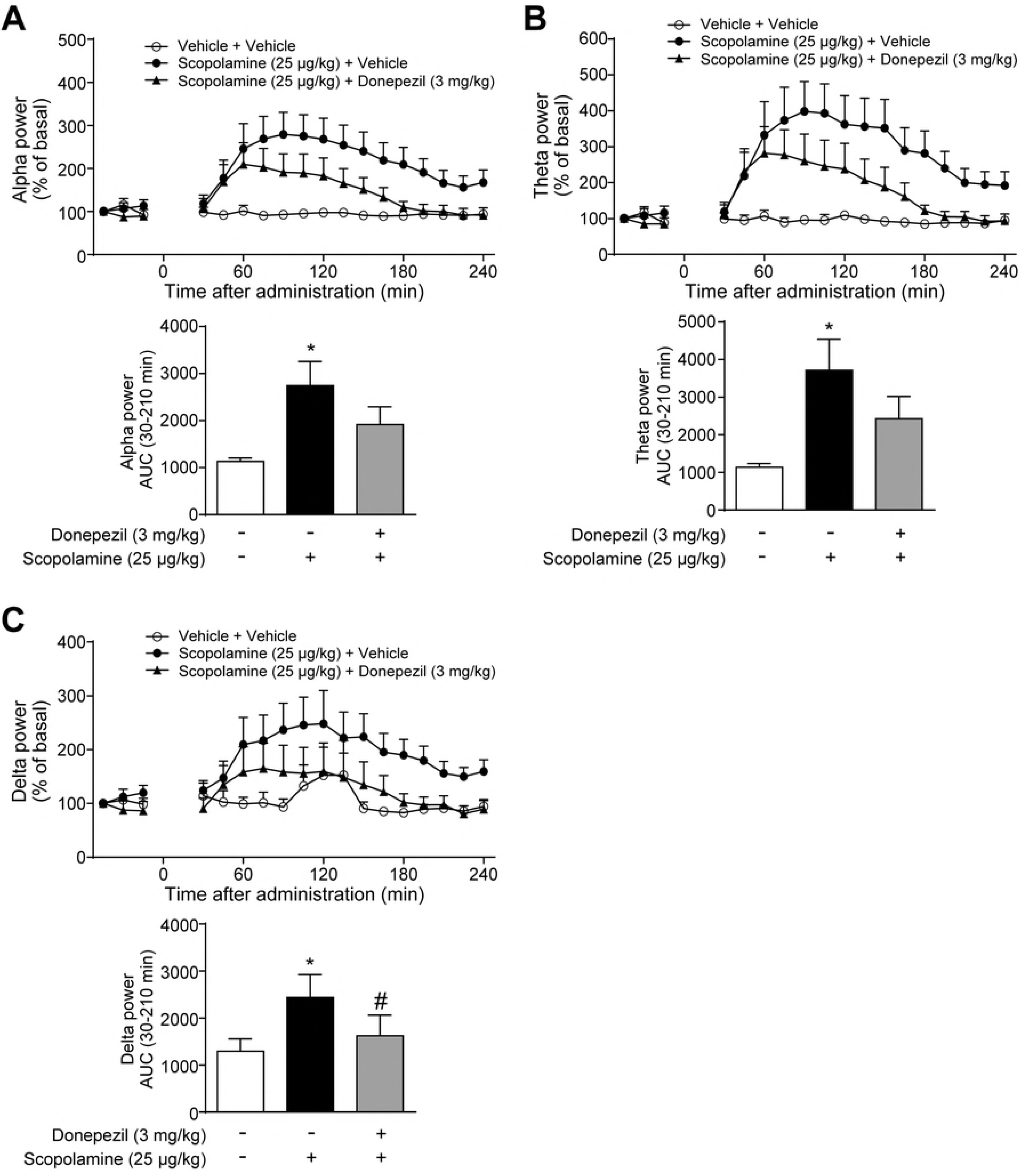
Donepezil at 3 mg/kg inhibited the scopolamine-induced qEEG spectral changes. Scopolamine (25 µg/kg, s.c.) and donepezil (3 mg/kg, p.o.) were simultaneously administered to cynomolgus monkeys. After treatment with drugs or vehicle, (A) alpha, (B) theta, and (C) delta power bands of qEEG spectra were measured from 30 to 240 min. Results represent mean ± SEM of AUC between 30 and 210 min after treatment for 4 monkeys in each group. ^*^*P* ≤ 0.05 versus vehicle-treated group by paired *t*-test. ^#^*P* ≤ 0.05 versus scopolamine-treated group by paired *t*-test. AUC, area under the curve; p.o., oral administration; qEEG, quantitative electroencephalogram; SEM, standard error of the mean; s.c., subcutaneous administration.

Furthermore, we examined the effects of donepezil on qEEG power spectra. Oral treatment of donepezil at 0.3 mg/kg alone increased alpha power band, whereas the dose-dependency was not observed (S3 Fig A). Donepezil (0.3 and 3 mg/kg) did not affect theta and delta power bands (S3 Fig B and C).

### Xanomeline, an orthosteric M_1_/M_4_R agonist, tended to attenuate the scopolamine-induced increases in qEEG power bands

Donepezil non-selectively activates both nicotinic and muscarinic receptors by increasing ACh levels in the synaptic clefts, thus, we next assessed the relative contribution of muscarinic receptors, especially M_1_R and M_4_R, to the effects of donepezil on scopolamine-induced qEEG power spectral changes, using an orthosteric M_1_/M_4_R agonist, xanomeline. The dose of xanomeline was determined based on the previous report, which demonstrated that subcutaneous treatment of xanomeline at 1 mg/kg exhibited antipsychotic-like activity without inducing sedative effects in monkeys [34]. Simultaneous administration of xanomeline (1 mg/kg, s.c.) and scopolamine (25 µg/kg, s.c.) showed a trend to suppress the effects of scopolamine on qEEG power spectral changes (Fig 3A–C). The scopolamine-induced increases in alpha, theta, and delta power bands were continuously inhibited between 60 and 240 min after treatment, although these effects were not statistically significant (Fig 3A–C). Thus, M_1_R and/or M_4_R antagonism may contribute to the qEEG power spectral changes caused by scopolamine treatment.

**Fig 3.**
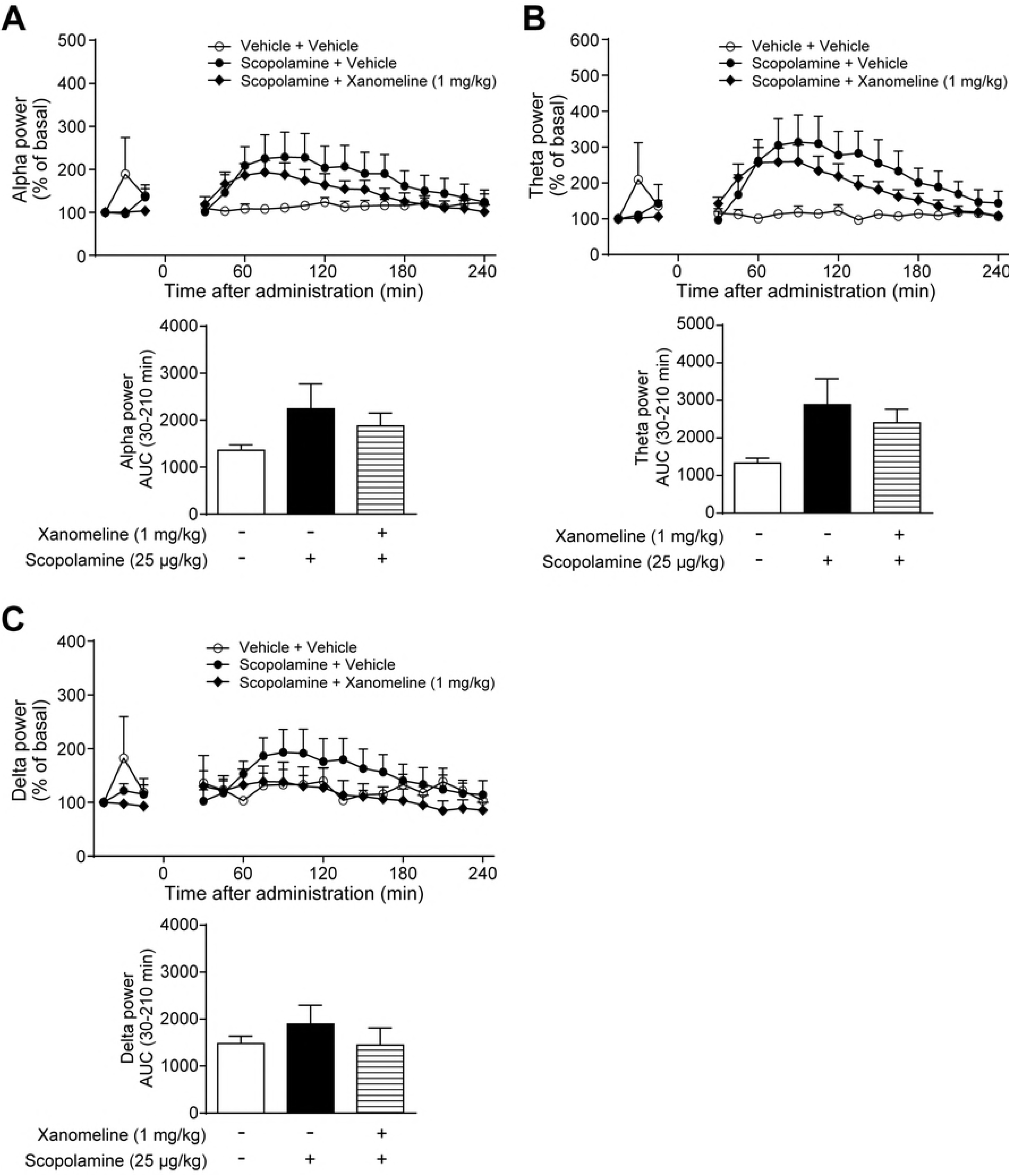
Xanomeline at 1 mg/kg tended to attenuate the scopolamine-induced qEEG changes. Scopolamine (25 µg/kg, s.c.) and xanomeline (1 mg/kg, s.c.) were simultaneously administered to cynomolgus monkeys. After treatment with drugs or vehicle, (A) alpha, (B) theta, and (C) delta power bands of qEEG spectra were measured from 30 to 240 min. Results represent mean ± SEM of AUC between 30 and 210 min after treatment for 4 monkeys in each group. AUC, area under the curve; qEEG, quantitative electroencephalogram; SEM, standard error of the mean; s.c., subcutaneous administration.

### TAK-071 inhibited the scopolamine-induced qEEG spectral changes in a dose-dependent manner

Based on the results of donepezil and xanomeline in combination with scopolamine, it was hypothesized that the scopolamine-induced qEEG power spectral changes may be useful translational biomarkers for the evaluation of TAK-071 effectiveness. Thus, we investigated whether TAK-071 affected the qEEG power spectral changes caused by scopolamine treatment. We previously reported that TAK-071 at 0.3 mg/kg significantly ameliorated the scopolamine-induced cognitive deficits, with a maximum plasma comcentration of 232 ng/mL in rats [11]. Based on the PK profile of TAK-071 in monkeys (S2 Table), the doses of TAK-071 were selected at 0.3–3 mg/kg to cover the effective plasma concentration in rats. Monkeys were simultaneously administered TAK-071 (0.3–3 mg/kg, p.o.) and scopolamine (25 µg/kg, s.c.). TAK-071 at 1 mg/kg significantly reduced the scopolamine-induced increase in alpha power band, whereas at 3 mg/kg, TAK-071 significantly suppressed the increases in both alpha and theta power bands (*P* ≤ 0.05, Fig 4A and B). Although there was no significant effect on delta power band, a decreasing trend was observed when TAK-071 was administered at 1 and 3 mg/kg (Fig 4C). Remarkably, these effects lasted between 90 and 240 min after drug administration (Fig 4A–C).

**Fig 4.**
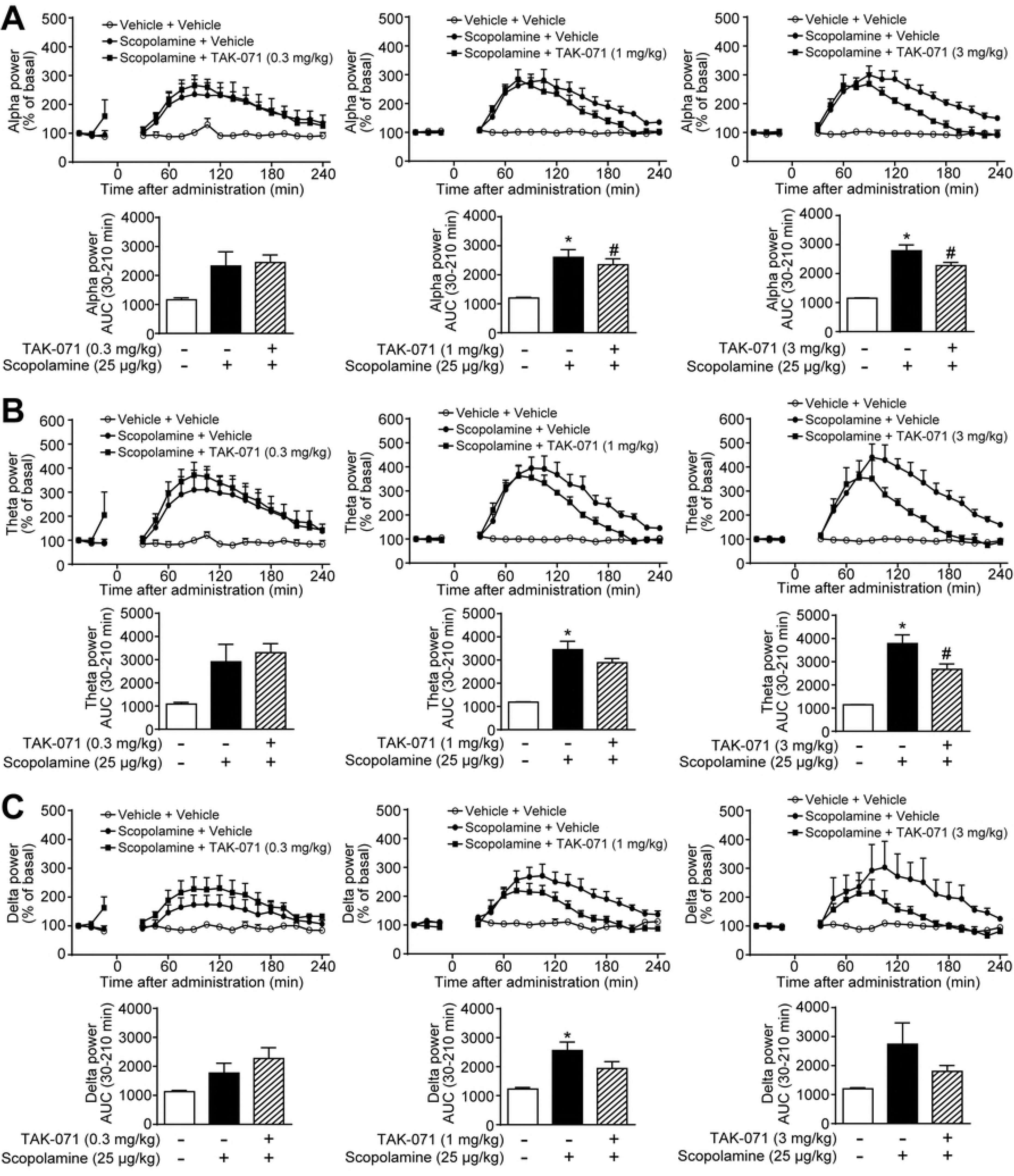
TAK-071 at 0.3–3 mg/kg suppressed the scopolamine-induced qEEG changes. Scopolamine (25 µg/kg, s.c.) and TAK-071 (0.3–3 mg/kg, p.o.) were simultaneously administered to cynomolgus monkeys. After treatment with drugs or vehicle, (A) alpha, (B) theta, and (C) delta power bands of qEEG spectra were measured from 30 to 240 min. Results represent mean ± SEM of AUC between 30 and 210 min after treatment for 3–4 monkeys in each group. ^*^*P* ≤ 0.05 versus vehicle-treated group by paired *t*-test. ^#^*P* ≤ 0.05 versus scopolamine-treated group by paired *t*-test. AUC, area under the curve; p.o., oral administration; qEEG, quantitative electroencephalogram; SEM, standard error of the mean; s.c., subcutaneous administration.

In addition to the scopolamine challenge paradigm, the effects of TAK-071 alone on qEEG power spectra were evaluated. Oral treatment of TAK-071 significantly decreased alpha and theta power bands at 1 and 3 mg/kg and at 1 mg/kg, respectively (*P* ≤ 0.05, S4 Fig A and B). In contrast, TAK-071 (0.3–3 mg/kg) did not change delta power band (S4 Fig C).

## Discussion

To improve the success rates in CNS drug development, appropriate translational biomarkers from preclinical to clinical stages are required. Neuroimaging techniques and qEEG measures are now being used to provide evidence of drug penetration into the CNS and determine the PK/PD properties of drugs [35-40].

Typical EEG abnormalities in patients with AD and DLB are associated with a decrease in alpha power band, which leads to increases in theta and delta power activities [16-19]. Several reports have demonstrated that scopolamine caused changes in qEEG power spectra in healthy subjects [27, 30, 31], which in part corresponded to those in patients with AD and DLB. Therefore, it is considered that the scopolamine model could reflect some aspects of the cholinergic deficit in AD and DLB in both PD and cognition. In addition to the findings from clinical studies, a scopolamine-induced increase in delta power band was also detected in freely moving rats [28, 29]. In line with the findings of clinical and preclinical studies, in this study we showed that scopolamine caused qEEG spectral changes in a dose-dependent manner, in particular, increases in theta and delta power bands in monkeys (Fig 1). We selected cynomolgus monkeys, not rodents, because monkey brains more closely resemble human brains than do rodents. Furthermore, monkeys are diurnal animals, and we considered it essential that the studies be performed under the same conditions as clinical studies, i.e., during daytime.

Among the scopolamine-induced changes in qEEG power spectra, the increases in theta and delta power bands in monkeys were similar to those reported in humans. Therefore, at least the effects of cholinergic agents in these power bands could be useful as translational biomarkers in the scopolamine challenge paradigm.

On the other hand, the increase in alpha power band in monkeys (Fig 1A) was opposite to the results in humans [27, 31]. It is known that alpha activity of the EEG power spectra is enhanced during an eyes-closed resting period and suppressed by visual stimulation in healthy individuals [41]. In clinical studies, subjects were instructed to remain awake and move as little as possible while keeping their eyes closed during qEEG measurements, so that the alpha power band would be increased under the measurement conditions. In contrast, the condition of the monkeys’ eyes during the qEEG measurements were uncontrollable. Monkeys used in our study would open their eyes because the experiments were conducted during the diurnal period under arousal conditions. This discrepancy might have led to the opposite effect on the alpha power band.

As shown in Fig 1, the time-dependent change of each qEEG spectrum during scopolamine treatment was relatively consistent with its PK profile (S1 Fig and S1 Table). Therefore, the current study suggests that the PK/PD relationship of scopolamine was detected in monkeys.

In line with the previous studies that reported significant suppression of the scopolamine-induced increase in delta power band by AChE inhibitors in rats [28, 29], this study also demonstrated that donepezil counteracted the scopolamine-induced increases in alpha, theta, and delta power bands (Fig 2). Furthermore, both xanomeline and TAK-071 showed the potential to inhibit the scopolamine-induced qEEG spectral changes (Figs 3 and 4). Taken together, these results suggest that the scopolamine-induced increases in alpha, theta, and delta power bands were suppressed partly through M_1_R activation in monkeys. In addition to the scopolamine challenge paradigm, we examined the effects of donepezil (0.3 and 3 mg/kg) and TAK-071 (0.3–3 mg/kg) on qEEG power spectra, and found that TAK-071 treatment alone slightly decreased alpha and theta power bands, whereas donepezil treatment alone did not decrease the qEEG power spectra (S3 and S4 Figs). Similar results were reported by Uslaner, where MK-7622, a selective M_1_ PAM, showed reductions of alpha and theta power bands [42]. The discrepancy between TAK-071 and donepezil may be explained by selective M_1_R signal transduction. In our previous study, TAK-071 was demonstrated to be a highly selective M_1_ PAM [11]. In contrast, donepezil indirectly and non-selectively activates both nicotinic and muscarinic receptors by increasing ACh levels in the synaptic clefts, which might lead to a masking effect on the decrease in qEEG power spectra through M_1_R activation. In fact, another AChE inhibitor, galantamine, did not change theta power band in healthy young volunteers [43].

In this study, we did not examine the direct association between qEEG spectral changes and cognitive deficits by scopolamine treatment in monkeys. Thus, it would be difficult to predict the effective dose of TAK-071 for cognitive improvements from the qEEG results in monkeys because the relationship between the effective doses at which cognitive improvement is observed and the induction of qEEG spectral changes remains unclear. However, both oral administration of 3 mg/kg donepezil and intramuscular administration of 0.1 mg/kg xanomeline improved scopolamine-induced cognitive deficits in monkeys [33, 44]. Furthermore, it is considered that qEEG abnormalities are correlated with the severity of cognitive impairment in AD and DLB [20-22]. While no direct comparisons were made in this study, inferences might be drawn about the effects on qEEG power spectra and cognition in monkeys. Hence, scopolamine-induced qEEG spectral changes may also be useful as a translational biomarker in humans for evaluation of procognitive effects of TAK-071.

## Acknowledgments

The authors thank Ms. Maki Miyamoto for performing the pharmacokinetic analysis of the test compounds, and Dr. Masami Yamada for providing TAK-071.

## Supporting information captions

**S1 Fig. Pharmacokinetic profiles of scopolamine after subcutaneous administration in cynomolgus monkeys.** Scopolamine (10 or 20 µg/kg) was administered subcutaneously to cynomolgus monkeys. After treatment with scopolamine, plasma samples were collected at 5, 15, 30, 60, 120 and 240 min. Results represent mean ± SD for 3 monkeys in each group.

SD, standard deviation.

**S2 Fig. Scopolamine did not change gamma and beta power bands of qEEG spectra in cynomolgus monkeys.** Scopolamine (25–100 µg/kg) was administered subcutaneously to cynomolgus monkeys. After treatment with scopolamine or vehicle, (A) gamma and (B) beta power bands of qEEG spectra were measured from 30 to 240 min. Results represent mean ± SEM of AUC between 30 and 210 min after treatment for 4 monkeys in each group.

AUC, area under the curve; qEEG, quantitative electroencephalogram; SEM, standard error of the mean.

**S3 Fig. Donepezil (0.3 and 3 mg/kg) did not affect qEEG power spectra in cynomolgus monkeys.** Donepezil (0.3 or 3 mg/kg, p.o.) was administered to cynomolgus monkeys. After treatment with donepezil or vehicle, (A) alpha, (B) theta, and (C) delta power bands of qEEG spectra were measured from 30 to 240 min. Results represent mean ± SEM of AUC between 30 and 210 min after treatment for 3–4 monkeys in each group. ^*^*P* ≤ 0.05 versus vehicle-treated group by paired *t*-test.

AUC, area under the curve; p.o., oral administration; qEEG, quantitative electroencephalogram; SEM, standard error of the mean.

**S4 Fig. TAK-071 (0.3–3 mg/kg) slightly, but significantly, decreased alpha and theta power bands of qEEG spectra in cynomolgus monkeys. TAK-071 (0.3**–3 mg/kg, p.o.) was administered to cynomolgus monkeys. After treatment with TAK-071 or vehicle, (A) alpha, (B) theta, and (C) delta power bands of qEEG spectra were measured from 30 to 240 min. Results represent mean ± SEM of AUC between 30 and 210 min after treatment for 4 monkeys in each group. ^*^*P* ≤ 0.05 versus vehicle-treated group by paired *t*-test.

AUC, area under the curve; p.o., oral administration; qEEG, quantitative electroencephalogram; SEM, standard error of the mean.

**S1 Table. Pharmacokinetic profiles of scopolamine after subcutaneous administration in cynomolgus monkeys.** Scopolamine (10 or 20 µg/kg) was administered subcutaneously to cynomolgus monkeys. After treatment with scopolamine, plasma sample was collected at 5, 15, 30, 60, 120, and 240 min. C_max_, T_max_ and AUC_0_–_4h_ were calculated from the results of S2 Fig. Results represent mean ± SD for 3 monkeys in each group.

**S2 Table. Pharmacokinetic profiles of TAK-071 after oral administration in cynomolgus monkeys.** TAK-071 (0.1 or 1 mg/kg) was orally administered to cynomolgus monkeys. After treatment with TAK-071, plasma sample was collected at 15, 30 min, and 1, 2, 4, 6, 8, 24, 48, and 72 hr. C_max_, T_max_ and AUC_0_–_72h_ were determined. Results represent mean ± SD for 3 monkeys in each group.

